# PPalign: Optimal alignment of Potts models representing proteins with direct coupling information

**DOI:** 10.1101/2020.12.01.406504

**Authors:** Hugo Talibart, François Coste

## Abstract

**Background:** To assign structural and functional annotations to the ever increasing amount of sequenced proteins, the main approach relies on sequence-based homology search methods, e.g. BLAST or the current state-of-the-art methods based on profile Hidden Markov Models (pHMM), which rely on significant alignments of query sequences to annotated proteins or protein families. While powerful, these approaches do not take coevolution between residues into account. Taking advantage of recent advances in the field of contact prediction, we propose here to represent proteins by Potts models, which model direct couplings between positions in addition to positional composition, and to compare proteins by aligning these models. Due to non-local dependencies, the problem of aligning Potts models is hard and remains the main computational bottleneck for their use.

**Results:** We introduce here an Integer Linear Programming formulation of the problem and PPalign, a program based on this formulation, to compute the optimal pairwise alignment of Potts models representing proteins in tractable time. The approach is assessed with respect to a non-redundant set of reference pairwise sequence alignments from SISYPHUS benchmark which have lowest sequence identity (between 3% and 20%) and enable to build reliable Potts models for each sequence to be aligned. This experimentation confirms that Potts models can be aligned in reasonable time (1′37″ in average on these alignments). The contribution of couplings is evaluated in comparison with HHalign and PPalign without couplings. Although Potts models were not fully optimized for alignment purposes and simple gap scores were used, PPalign yields a better mean *F*_1_ score and finds significantly better alignments than HHalign and PPalign without couplings in some cases.

**Conclusions:** These results show that pairwise couplings from protein Potts models can be used to improve the alignment of remotely related protein sequences in tractable time. Our experimentation suggests yet that new research on the inference of Potts models is now needed to make them more comparable and suitable for homology search. We think that PPalign’s guaranteed optimality will be a powerful asset to perform unbiased investigations in this direction.

## Background

Thanks to sequencing technologies, the number of available protein sequences has considerably increased in the past years, but their functional and structural annotation remains a bottleneck. This task is thus classically performed *in silico* by scoring the alignment of new sequences to well-annotated homologs. One of the best-known method is BLAST [1], which performs pairwise sequence alignments. The main tools for homology search are now based on profile Hidden Markov Models (pHMMs), which model position-specific composition, insertion and deletion probabilities of each family of homologous proteins. Two well-known software packages using pHMMs are widely used today: HMMER [2] aligns sequences to pHMMs and HH-suite [3] takes it further by aligning pHMMs to pHMMs.

Despite their solid performance, pHMMs are innerly limited by their positional nature. Yet, it is well-known that residues that are distant in the sequence can interact and co-evolve, e.g. due to their spatial proximity, resulting in correlated positions. One can cite for instance experiments of Ranganathan et al. on the WW domain who showed by experimentally testing libraries of artificial sequences of the WW domain that coevolution information is necessary to reproduce the functional properties of native proteins [4].

There have been a few attempts to make use of long-distance information. Menke, Berger and Cowen introduced a Markov Random Field (MRF) approach, SMURF [5], where MRFs generalize pHMMs by allowing dependencies between paired residues in *β*-strands to recognize proteins that fold into *β*-structural motifs. Their MRFs are trained on multiple structure alignments. A model simplification [6] and heuristics [7] have been proposed to speed up the process. While these methods outperform HMMER[2] in propeller fold prediction, they are limited to sequence-MRF alignment on *β*-strand motifs with available structures. Xu et al. [8] proposed a more general method, MRFalign, which performs MRF-MRF alignments using probabilities estimated by neural networks from amino acid frequencies and mutual information. Unlike SMURF, MRFalign handles dependencies between all positions and MRFs are built from multiple sequence alignments. In addition to these inputs, MRFalign relies on complex scoring functions based on Conditional Neural Fields and Probabilistic Neural Network trained on reference alignments and structural information to optimize the similarity measures of the positional and coupling potentials of the MRF models to be compared. In reported results, PSSM-PSSM and HMM-HMM alignment methods are outperformed by MRFalign in terms of both alignment accuracy and remote homology detection accuracy, notably on mainly beta proteins, showing the potential of using long-distance information in protein sequence alignment.

Meanwhile, a more interpretable type of MRF grounded in the maximum entropy principle led to a breakthrough in the field of contact prediction [9]: the Potts model. This model was brought forward by Direct Coupling Analysis [10], a statistical method to extract direct correlations from multiple sequence alignments. Once inferred on a multiple sequence alignment (MSA), a Potts model’s nodes represent positional conservation, and its edges represent direct couplings between positions in the MSA. Unlike mutual information which also captures indirect correlations between positions, Potts models are global models capturing the collective effects of entire networks of correlations through their coupling parameters [11], thus tackling indirect effects and making them a relevant means of predicting interactions between residues. Beyond contact prediction, the positional and the direct coupling information captured by Potts model’s parameters might also be valuable in the context of protein homology search. The idea of using Potts models for this purpose was proposed last year at the same workshop by Muntoni and Weigt [12], proposing to align sequences to Potts models, and by us [13], proposing to align Potts models to Potts models in our generic framework for the comparison of protein sequences using direct coupling information named ComPotts.

The main computational bottleneck for such approaches is that, due to non-local dependencies, alignment problems involving Potts models are hard. Muntoni and Weigt [12] proposed an approximate message-passing algorithm to align a sequence to a Potts model. In this paper, we fully describe PPalign, our method introduced in ComPotts to optimally align two Potts models representing proteins in tractable time and focus on its performances in terms of alignment quality on remote homologs. In the following sections, we explain our choices for the inference of Potts models and describe the method for aligning them, which builds on the work of Wohlers, Andonov, Malod-Dognin and Klau [14, 15, 16] to propose an Integer Linear Programming formulation for this problem, with an adequate scoring function. To assess the tractability and the quality of PPalign’s alignments, we extracted 33 non-redundant pairwise reference alignments with a particularly low identity from the manually curated structural alignments database SISYPHUS [17] and randomly split it into a training set of 11 pairs to train our hyperparameters and a test set of 22 pairs on which we compared our results with HHalign’s alignments of pHMMs built on the same input data. On this test set, our method yielded the exact solutions up to a chosen epsilon in tractable time, and outperformed HHalign in terms of alignment quality with an *F*_1_ score better on average and significantly better for 5 alignments, suggesting that direct couplings can improve alignment quality of remote homologs.

## Methods

### Inference of Potts models

Potts models are discrete instances of pairwise Markov Random Fields which originate from statistical physics. They generalize Ising models by describing interacting spins on a crystalline lattice with a finite alphabet. In the paper introducing Direct Coupling Analysis, Weigt et al. came up with the idea of applying them to proteins: by building a multiple sequence alignment of a protein sequence and its close homologs and inferring a Potts model on it, one can predict contacts between residues by looking at its parameters [10].

The inference of a Potts model from a set of protein sequences can be formally defined as follows:

Let *S* = {*s^n^*}_*n* = 1,⋯ *N*_ be a set of *N* protein sequences of lengths *l*_1_, ⋯, *l_N_*. A multiple sequence alignment (MSA) of these sequences can be defined as a set of *N* sequences *X* = {*x^n^*}_*n*=1,⋯, *N*_ on the alphabet of *S* extended with a new gap character ‘–’, which all have the same length *L* and such that removing all gaps from a sequence *x^n^* gives *s^n^*. By extension, L is called the length of the MSA. We denote by *q* the size of the alphabet.

A Potts model with *q* states for MSA *X* can be defined as a statistical model whose probability distribution *P* over all sequences of length L maximizes the Shannon entropy *H*(*P*) = − ∑_*y* ∈ |1,⋯, *q*}^*L*^_ *P*(*y*) log *P*(*y*) and generates the empirical single and double frequencies of the MSA as marginals:

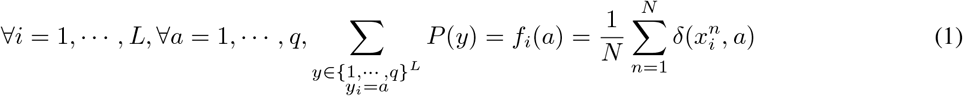

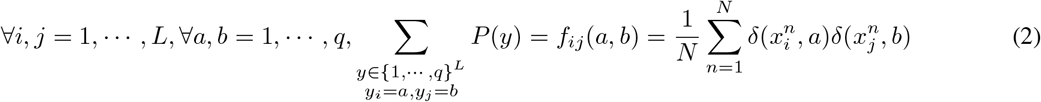

This probability distribution is unique and has the following form:

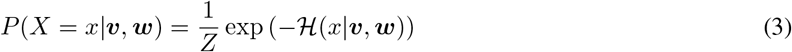

where *Z* is a normalization constant: 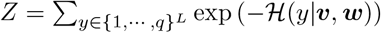 and 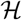 is an energy function defined as

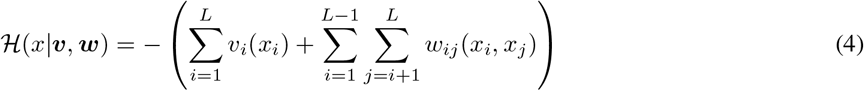

where the parameters (***v, w***) that define a Potts model are the ones that maximize the likelihood of the sequences in the MSA *X*:

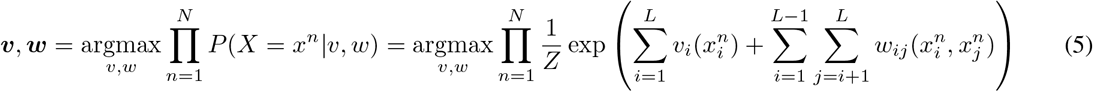

These parameters can be assigned a practical interpretation:

- ***v*** = {*v_i_*}_*i*=1,⋯, *L*_ are positional parameters termed “fields”. Each *v_i_* is a real vector of length *q* where *v_i_*(*a*) is related to the propensity of letter *a* to be found at position *i*.
- ***w*** = {*w_ij_*}_*i,j*=1,⋯, *L*_ are pairwise coupling parameters. Each *w_ij_* is a *q × q* real matrix where *w_ij_*(*a, b*) quantifies how compatible letters *a* and *b* are when found at positions *i* and *j*.

An illustration of Potts model is given Figure 1.

**Figure 1:**
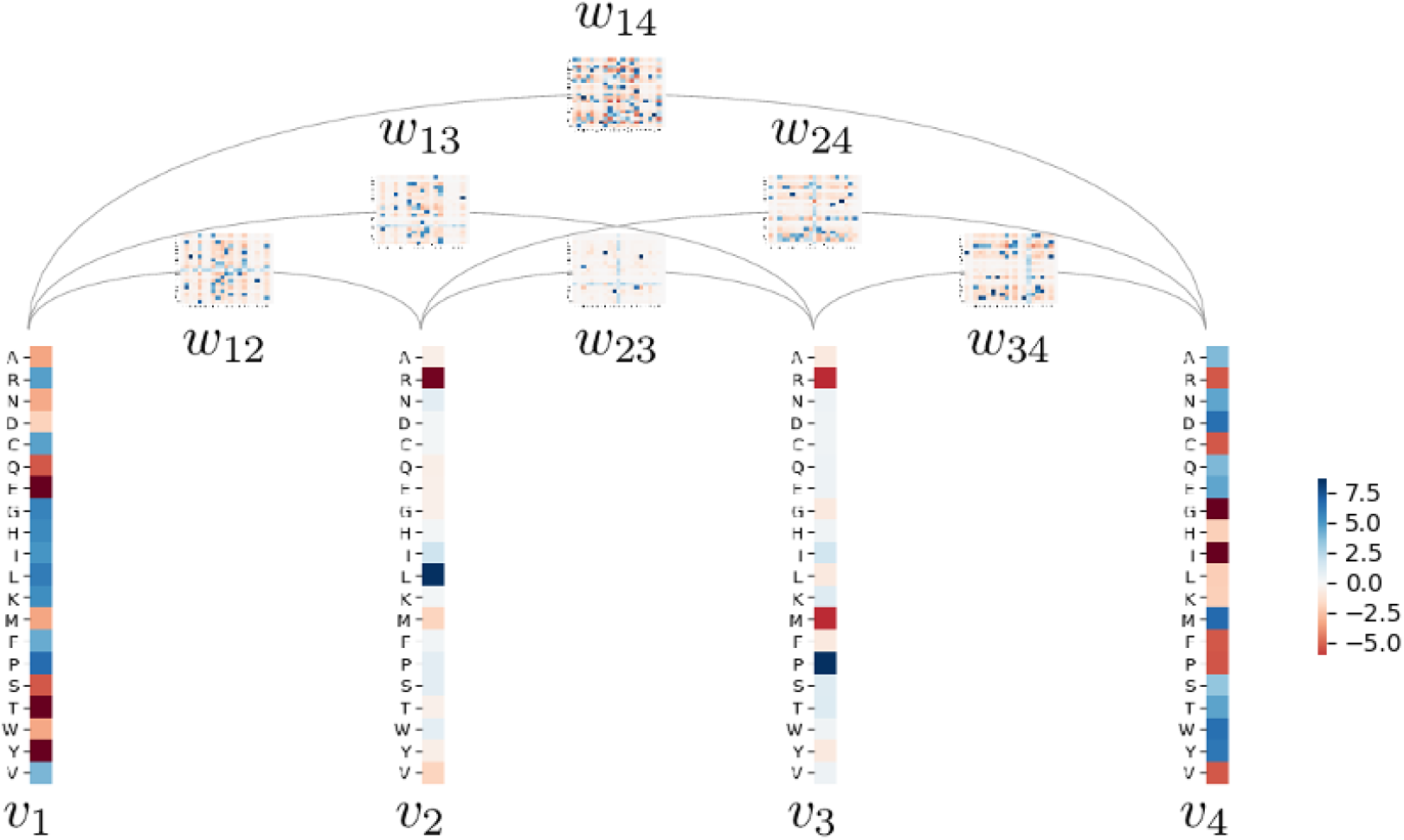
Example of Potts model representing a MSA of length 4. Each column in the MSA is associated with a field vector *v_i_* of length *q* = 20 where each *v_i_*(*a*) is a real value weighting positively or negatively the occurrence of letter *a* at position *i*. Each pair of positions (*i, j*) is associated with a *q × q* coupling matrix *w_ij_* where *w_ij_*(*a, b*) are real values weigthing positively or negatively the co-occurrence of letters *a* and b respectively at position *i* and *j*.

In practice, maximizing the likelihood would require the computation of the normalization constant *Z* at each step, which is computationally intractable. Among the several approximate inference methods that have been proposed [18, 19, 20, 21, 11], we opted here for pseudo-likelihood maximization since it was proven to be a consistent estimator in the limit of infinite data [22, 23] within reasonable time. Furthermore, since our goal is to align Potts models, we need the inferrence to be geared towards similar models for similar MSAs, which is not what inference methods were initially designed for. In an effort towards inferring canonical Potts models, we have chosen here to use CCMpredPy [24], a recent Python-based version of CCMpred [25] which, instead of using the standard *L*_2_ regularization prior 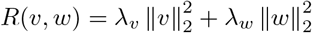, allows us to use a smarter prior on *v*:

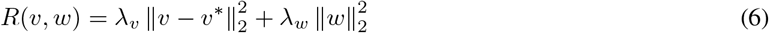

where *v*^*^ obeys

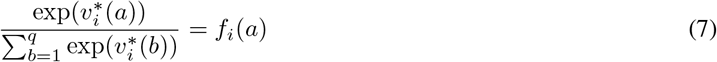

which yields the correct probability model if no columns are coupled, i.e. 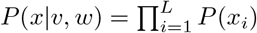. Our intuition is that positional parameters should explain the MSA as much as possible and only necessary couplings should be added.

### From a protein sequence to a Potts model

To add coupling information to a protein sequence, the first step is to build a MSA of its close homologs to get sufficient coevolutionary signal. In this paper, based on CCMpred’s recommendations [26], for each sequence we run HHblits [3] v3.03 with the following parameters:

`-maxfilt 100000 -realign_max 100000 -all -B 100000 -Z 100000 -n 3 -e 0.001` on Uniclust30 [27] (08/2018 release), and then process the output by:

- filtering at 80% identity using HHfilter
- taking the first 1000 sequences
- removing all columns with > 50% gaps using trimal [28]

The resulting MSA is inputted to CCMpredy [24] using default parameters to infer a Potts model, and trimmed positions *i* (with > 50% gaps in the input MSA) are re-inserted in the model with positional parameters at position *i* set to background fields defined using frequencies *f*_0_ given by [29]

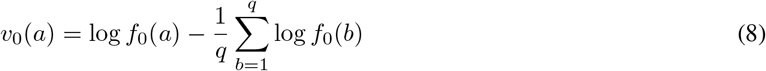

and pairwise coupling parameters with position *i* set to:

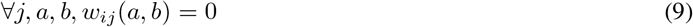

### Parameter rescaling strategy

Since existing Potts model inference methods were specifically designed for the prediction of co-evolving position pairs, inferred parameters might not be ideally suited for Potts model comparison. This section describes two strategies implemented to compensate for these shortcomings.

### Lessening the effect of small sample variations on the positional parameters

Since field parameters v are linked to single frequencies through a logarithmic relation (see equation (7)), any noise in the presence of small probabilities can have a great impact on the model parameters. This has a dramatic effect on the scoring function we use for pairwise Potts model alignment since the sign of each parameter directly determines the sign of their similarity score (see next section). To lessen the effects of sampling variations, we apply additive smoothing to the softmax probability distribution *p_i_* associated with each *v_i_*.

More formally, a standard softmax probability distribution *p_i_* is extracted for each positional parameter *v_i_*:

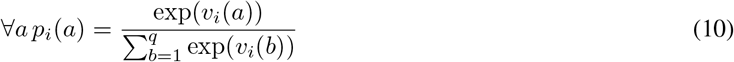

It is then smoothed towards a uniform distribution so that very low probabilities are more homogenized:

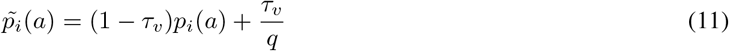

where *τ_v_* is a parameter controlling the amount of additive smoothing used. Final smoothed parameters 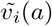 are retrieved by inverting the softmax function using the fact that 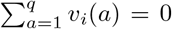 according to CCMpredPy’s gauge choice:

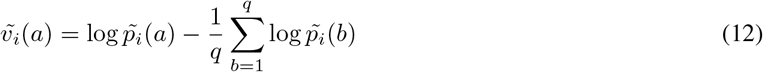

Summing up in one formula, each parameter *v_i_*(*a*) of the inferred Potts model is smoothed using the following function:

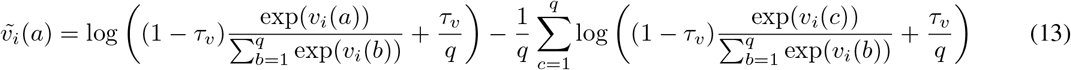

### Diminishing contributions of anti-correlations

In theory, coupling values inside a *w_ij_* matrix are supposed to deviate positively or negatively from 0 to reflect a (direct) correlation or anti-correlation. In practice however, while input data can be sufficient to assert that two letters a and b are likely to be found together at positions *i* and *j*, deducing that they should not be found together at positions *i* and *j* requires more examples to have sufficient countings on all pairs of *a* and *b*. Considering that our data set is limited, a large number of spurious anti-correlations can arise from a mere lack of data.

Since positive correlations are more likely to be supported by available training sample than negative ones, our approach here is to skew the coupling value distribution inside each *w_ij_* matrix to favor higher, positive values.

To do this, we extract each coupling matrix probability distribution as for the fields, only with a different softmax base *β_w_*, chosen so that the extracted distribution is skewed towards higher probabilities:

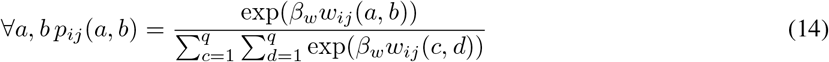

and, as for the fields, smooth it towards a uniform distribution to lessen noise, which gives:

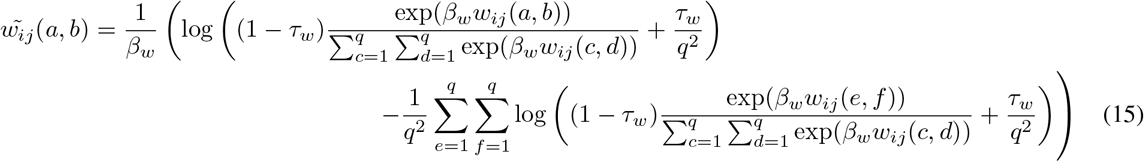

Using this smoothing scheme on each input Potts model make them more comparable since the most significant information stands out while sampling variations are tuned down.

### Alignment of Potts models

This section introduces our method for aligning two Potts models. The function we designed to score a given alignment is described and constraints ensuring that the alignment is proper are added as in Wohlers et al. [16], resulting in an Integer Linear Programming formulation that can be optimized using their efficient solver.

### Scoring function

Basically, the best alignment between two Potts models *A* = (***v**^A^, **w**^A^*) and *B* = (***v**^B^, **w**^B^*) of lengths *L_A_* and *L_B_* is defined as the alignment which maximizes the similarity between aligned fields and aligned couplings. Formally, this means finding the values of the binary variables *x_ik_* where *x_ik_* = 1 iff position *i* in Potts model *A* is aligned with position *k* in Potts model *B* so as to maximize:

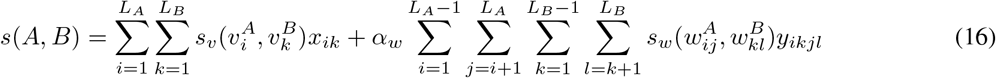

where 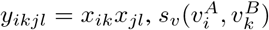 and 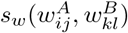 are similarity scores, respectively between positional parameters 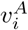 and 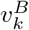 and coupling parameters 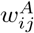 and 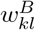 and *α_w_* is a coefficient ensuring proper balance between positional and coupling score.

To measure the similarity between vectors, the scalar product is a natural candidate. We propose thus to measure the similarity 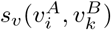 between field parameters using:

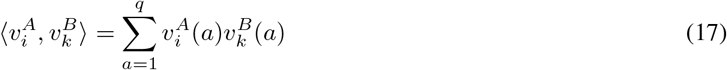

and to measure the similarity 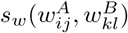 between coupling parameters by the extension of the scalar product to matrices, the Frobenius inner product:

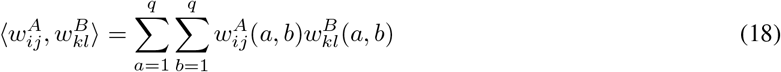

Note that this scoring function for two Potts models naturally generalizes the score of a sequence *x* for a given Potts model since its energy can be computed as:

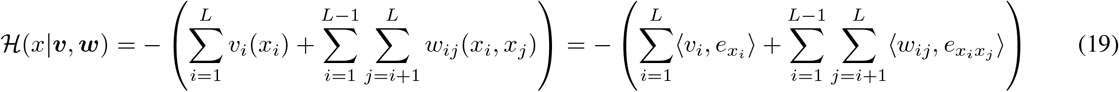

where:

- *e_x_i__* is the vector defined by ∀_*a*_ ∈ [1..*q*]; *e_x_i__* (*a*) = *δ*(*a, x_i_*)
- *e_x_i_x_j__* is the matrix defined by ∀(*a, b*) 2 [1..q]2; *e_x_i_x_j__* (*a, b*) = *δ*(*a, x_i_*)*δ*(*b, x_j_*)

Inspired by sequence alignment methods which use log-odds ratios to compute their scores with respect to a background model, we remove the background field *v*_0_ defined in equation (8) to each field vector before computing the scalar product. The actual similarity score between two positional parameters 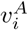 and 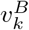 used in this paper is thus:

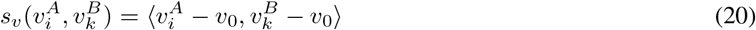

while the similarity score between two coupling parameters 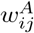 and 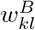 remains:

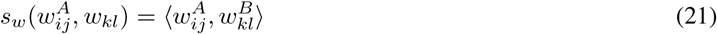

### Optimizing score with respect to constraints

Naturally, the scoring function should be maximized with respect to constraints ensuring that the alignment is proper. In that perspective, we build on the work of Wohlers et al. [16], initially dedicated to protein structure alignment, to propose an Integer Linear Programming formulation for the Potts model alignment problem.

Let us first introduce necessary definitions and notations following [16] to define a proper alignment.

The *alignment graph* of two Potts models *A* and *B* of lengths *L_A_* and *L_B_* is a *L_A_ × L_B_* grid graph where rows (from bottom to top) represent positions in *A* and columns (from left to right) represent positions in *B*. A node *i.k* in the alignment graph represents the alignment of node *i* from Potts model *A* and node *k* from Potts model *B*. Directed edges (*i.k, j.l*) are drawn for *i < j* and *k < l*. In this framework, an alignment of *n* positions in the two Potts models is represented by a set of nodes {*i*_1_.*k*_1_,⋯, *i_n_.k_n_*} where *i_n_* < ⋯ < *i_n_* and *k_n_* < ⋯ < *k_n_*, termed *increasing path*.

In order to properly set constraints on the alignment, two additional node sets are defined: row_*ik*_(*j*) (resp. col_*ik*_(*l*)) is the maximal set of nodes in the alignment graph that are tails of edges with head at *i.k* or heads of edges with tail at *i.k*, that contain at least one node at row *j* (resp. column *l*), and that *mutually contradict*, i.e. no two of them lie on an increasing path.

To cast the alignment problem into an ILP, binary variables *x_ik_* are assigned to each node *i.k* in the alignment graph, with *x_ik_* = 1 if position *i* in Potts model *A* and position *k* in Potts model *B* are aligned, and similarly a binary variable *y_ikjl_* is assigned to each edge in the alignment graph where *y_ikjl_* = 1 if edge (*i.k, j.l*) is activated.

Given notations above, the alignment of two Potts models *A* and *B* of lengths *L_A_* and *L_B_* and parameters (*v^A^, w^A^*), (*v^B^, w^B^*) can be formulated as the following Integer Linear Programming problem:

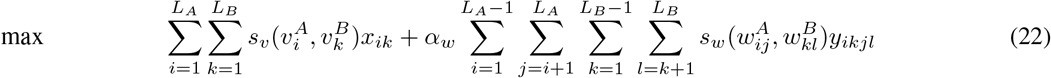

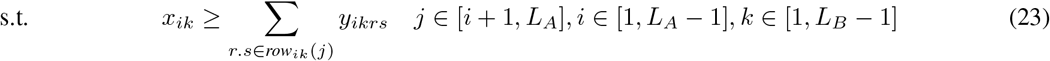

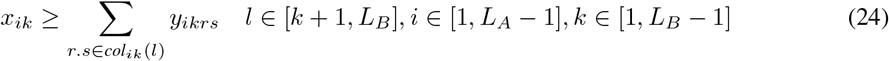

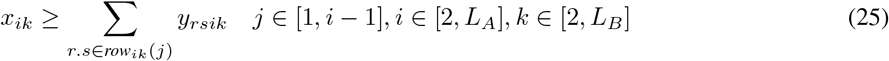

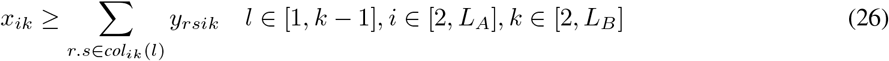

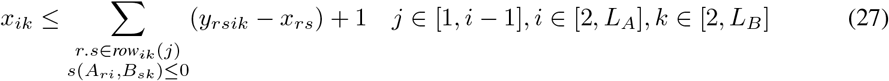

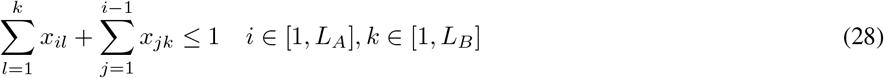

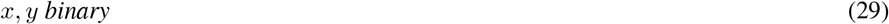

Constraints (23) and (24) prevent edges from activating if their tails are not activated and ensure that heads of edges with a common tail do not contradict, and constraints (25) and (26) denote the reverse situation. Constraint (27) ensures that edges are activated if their heads and tails are activated (this constraint is necessary since similarity scores can be negative). Finally, constraint (28) ensures that the nodes lie on an increasing path.

A major asset of the solver is that it can yield the exact solution of this ILP, or a solution within a chosen epsilon range of the exact one, in tractable time. Desired precision of the optimization can be set by the parameter *ϵ*, ensuring that 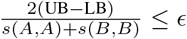 where UB and LB are the upper and lower bounds guaranteed by the solver for the solution, to avoid unnecessary optimization steps (the precision can be sufficient for the task) and speed up the search (often the last optimization steps only contribute to tighten the bounds while the optimal solution is already found).

### Gap cost and offset

As in [16], an affine gap cost function can be added to the score function to account for insertions and deletions in the sequences, with the appropriate choice of a gap open and a gap extend penalties.

Furthermore, as in most profile-profile methods [30], in order to prevent our method from greedily aligning every position, we penalize each aligned pair with a fixed negative offset hyperparameter.

### Data

To evaluate PPalign and the contribution of distant dependencies, we focused on reference alignments based on structures with low sequence identity. We opted for SISYPHUS database [17] since it provides manually curated structural alignments for proteins with non-trivial relationships. Our data set was built as follows:

- From each multiple sequence alignment in SISYPHUS, every possible pairwise sequence alignment with a sequence identity lower than 20% was extracted (we set a low sequence identity threshold to focus on harder targets)
- For each sequence in each of these extracted pairwise reference alignments, we attempted to build a Potts model with the workflow previously described. Sequences that had less than 1000 80% non-redundant homologs were discarded to focus on sequences with sufficient co-evolution signal. Due to CCMpredPy memory consumption, trimmed MSAs whose length was longer than 200 also had to be discarded.
- Finally, for each reference multiple sequence alignment in SISYPHUS with more than two of such eligible sequences, a reference sequence pair was randomly selected. This last steps discards many alignment pairs but ensures that no multiple sequence alignment biases the results.

This resulted in a set of 33 non-redundant reference pairwise alignments which was randomly split into a train set of 11 alignments on which our hyperparameters were trained (see table 1) and a test set of 22 target alignments (see table 2).

**Table 1:**
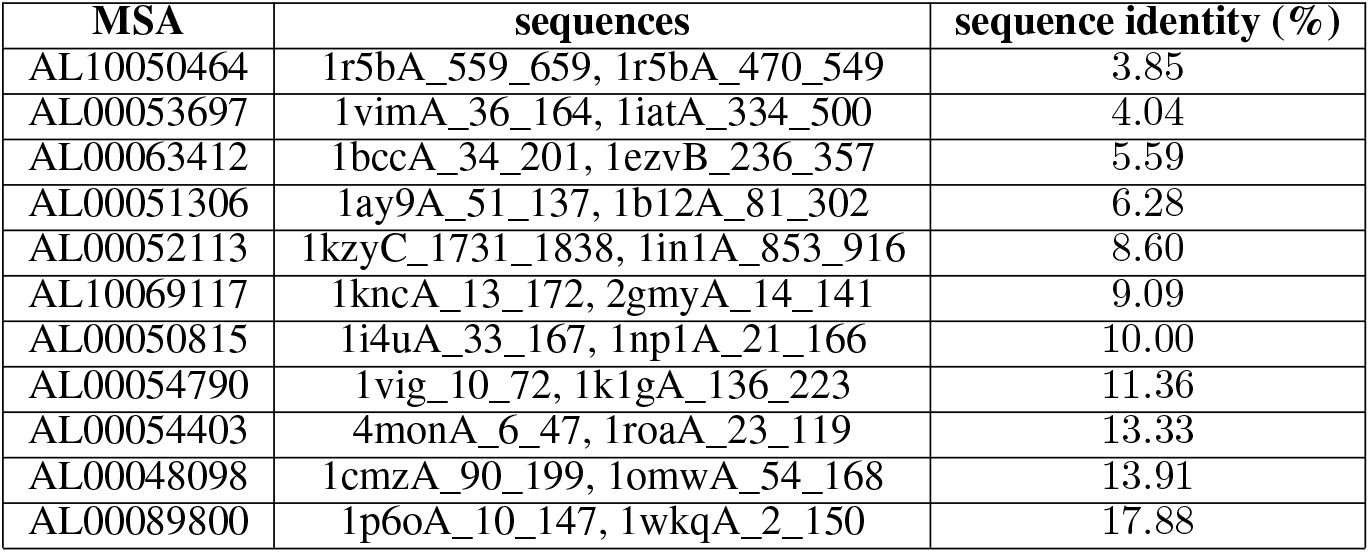
Training set.

**Table 2:**
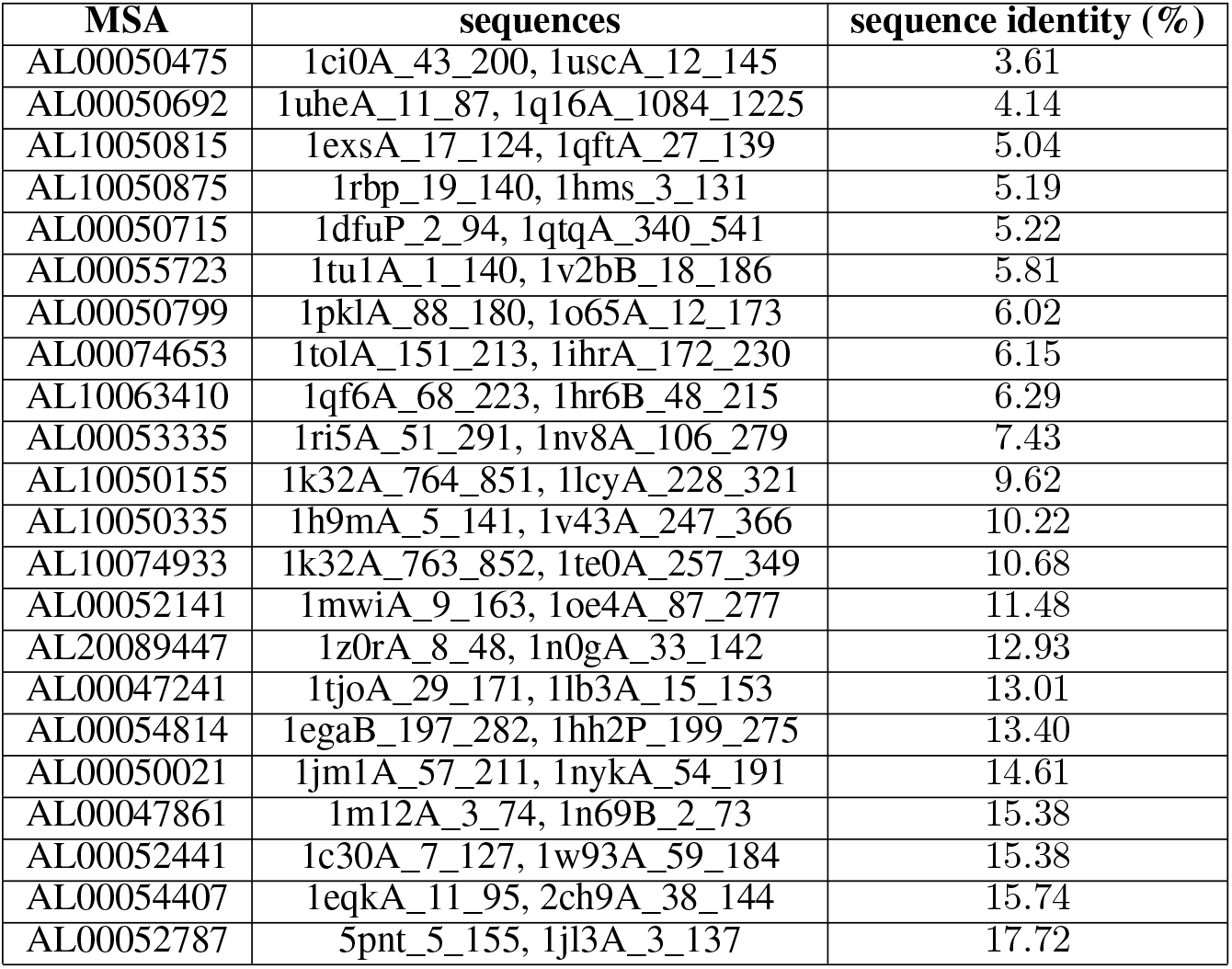
Test set.

### Alignment evaluation metrics

Alignment quality with respect to SISYPHUS’ reference alignments is assessed by computing alignment precision:

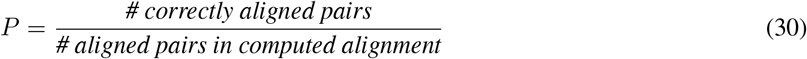

and recall:

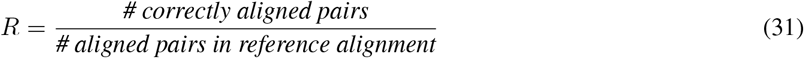

using Edgar’s qscore program [31] v2.1, and *F*_1_ score:

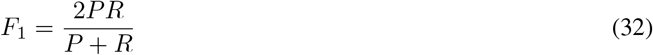

### PPalign’s hyperparameters

PPalign’s hyperparameters were optimized on the 11 alignments from the training set using Hyperopt library [32] to maximize the *F*_1_ score. This process showed to be excessively time-consuming, Hyperopt being unable to show a convergence on the choice of the parameters after one month. In order to reduce the hyperparameter search space and speed up the convergence of this process, we had to arbitrarily set some parameters after some trials on the training set: precision e was set to 0.02, *τ_v_* and *τ_w_* from equations (13) and (15) were both set to 0.4 and the gap extend penalty was set to 0. In accordance with the expected NP-hardness of the problem, time needed to find optimal alignment could be very long for some sets of parameters and even exceed the 6 hours time-out we set. We observed yet that good alignments were usually already found in less than 1 minute and decided to set the time-out by alignment to this value to speed-up more the optimisation of the remaining parameters by Hyperopt, which yielded the following values:

- Gap open penalty: 13
- kCoupling contribution coefficient *α_w_*: 6
- Softmaxbase *β_w_*: 8.0
- Offset *γ*: 1.0

### Other methods to be compared

In this experiment, we compared the results of PPalign with:

- PPalign without coupling score, i.e. *α_w_* = 0 (termed PPalign positional)
- HHalign v3.0.3, run with default options to align pHMMs built with HHmake with default options from the MSAs used to infer Potts models (except for the trimming of the positions with > 50% gaps since pHMMs handle well insertions and deletions)
- BLASTp v2.9.0+ without E-value cutoff, run on the sequences truncated as in our training MSAs, to provide an indication on the sequences’ similarity

## Results

### Tractable computation time

We examined the computation times of PPalign, PPalign positional and HHalign, considering the time they took to align the models (and not the steps to build them, that can be done offline) of the sequence pairs from the test set. Experiments were run on a Debian9 virtual machine with 4 VCPUs (2.3 GHz) and 8 GB RAM. The timeout for each alignment was set to 6 hours.

The first result is that all the alignments could be computed by PPalign in running times ranging from 5 seconds to 6 minutes, with an average of 1 min 36. Figure 2a plots the running times with respect to the lengths of the models to align. It shows that most problems (17/22) are easily solved and that running time for these problems increases gently with the lengths of the models, while a few (5/22) other problems stand out from this majority trend but are still solved in a few minutes.

**Figure 2:**
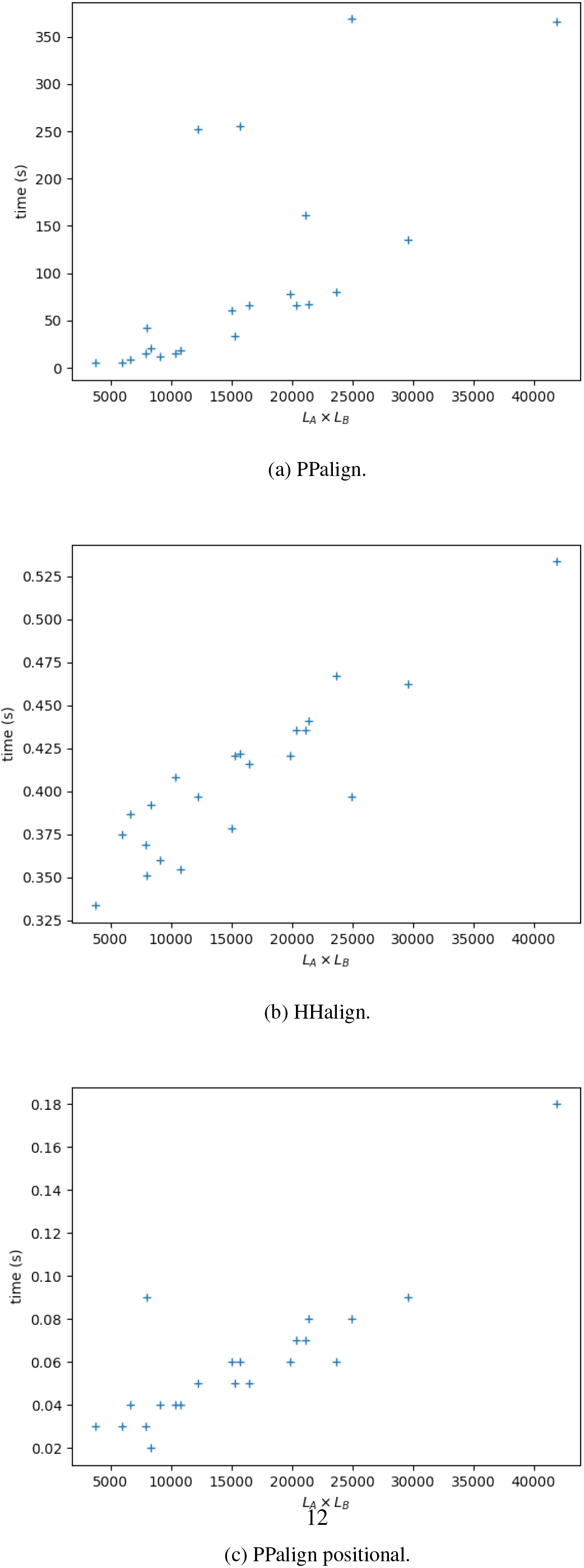
Time for aligning models of lengths *L_A_* and *L_B_* for sequence pairs from test set.

When couplings are not considered, the problem is fundamentally easier and running times of HHalign and PPalign positional are significantly faster than PPalign: both programs were able to compute each optimal positional alignment in less than 1 second. The running times of HHalign and PPalign positional are plotted in Figure 2b and Figure 2c. The two plots are not completely comparable since time needed to load the models is here included for HHalign and not for PPalign positional, but they illustrate the difference between the dynamic programming approach of HHalign, with a steady running time increment with the length of the models, and the Integer Linear Programming optimization approach of PPalign positional, showing here 2 outliers with respect to the general tendency.

### Alignment quality

Alignment quality was assessed by comparing the alignment obtained by the different methods for the 22 sequences pairs in the test set to their reference alignment.

Overall, PPalign achieves a better *F*_1_ score than HHalign (0.600 versus 0.578) with a better recall (0.613 vs 0.533) but a lower precision (0.587 vs 0.661), outperforming it in 12 out of the 22 alignments. BLAST only aligned 4 out of the 22 pairs, yielding an average *F*_1_ score of 0.113.

Results for each sequence pair of the test set are displayed in Figure 3.

**Figure 3:**
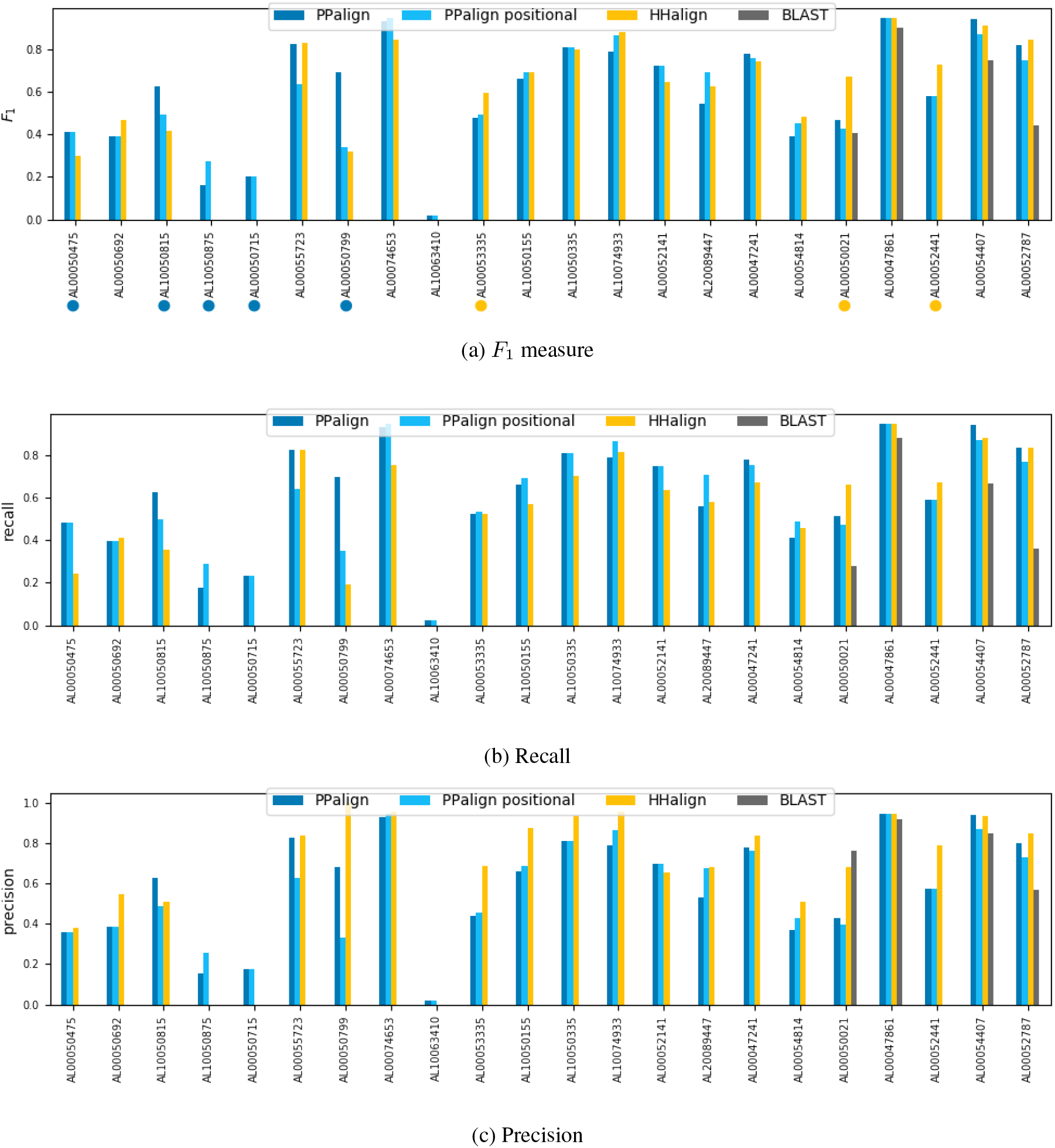
Quality of the alignments computed by PPalign, PPalign positional, HHalign and BLAST with respect to target reference alignments in test set (ordered by increasing percentage of sequence identity).

In most cases, PPalign and HHalign yield similar *F*_1_ scores (with less than 0.1 difference), except for 8 sequence pairs. 5 of them, marked by blue dots in the Figure 3a, are significantly better aligned by PPalign: AL00050475, AL00050692, AL10050875, AL00050715 and AL00050799 which are among the 7 alignments with the smallest percentage of sequence identity with respectively 3.61%, 5.04%, 5.19%, 5.22% and 6.02%. AL10050875 and AL00050715 are part with AL10063410 of the three sequence pairs that HHalign fails completely to align, yielding small and incorrect alignments with an *F*_1_ score of 0. On AL10063410, PPalign also failed, but on AL10050875 and AL00050715 it was able to do a bit better than HHalign by correctly aligning in each case roughly a fifth of the target alignment while still being wrong on the four other fifths. On AL00050475 and AL00050692, PPalign successfully retrieves about half of the target alignments when HHalign was retrieving only respectively a fifth and a third of it. The contribution of the coupling parameters is particularly noticeable for AL00050799, PPalign correctly retrieving almost 70% of the alignment while HHalign retrieves only 20% of it (see detailed analysis in Figure 4).

**Figure 4:**
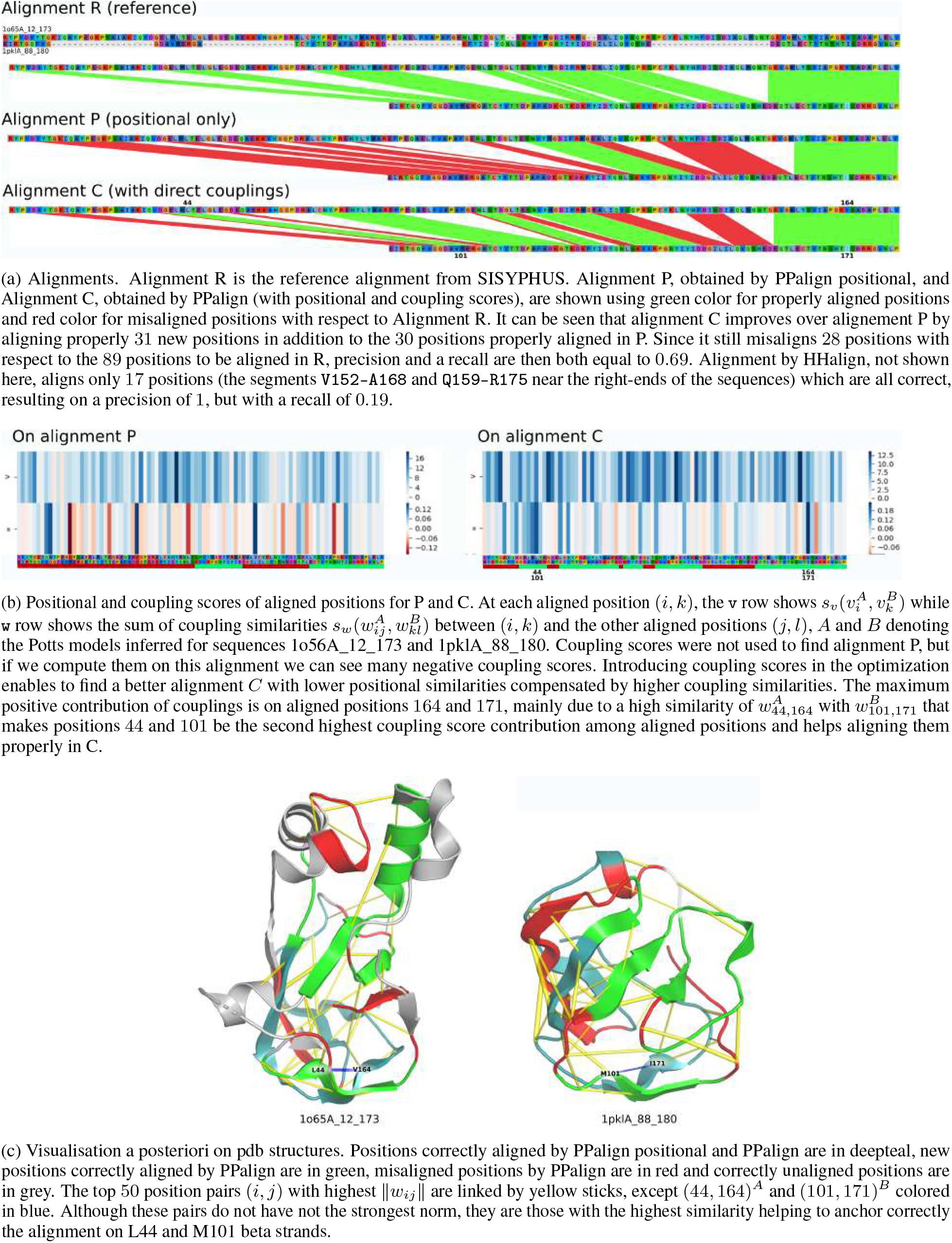
Illustration of the contribution of couplings for the alignment of 1o65A_12_173 and 1pklA_88_180 sequences.

PPalign is significantly outperformed by HHalign on 3 pairs, marked by yellow dots in Figure 3a. On AL00053335 (7.43% sequence identity), PPalign suffers from its tendency to align too many positions: like HHalign it correctly aligns half of the target alignment, but it proposes a longer alignment than HHalign, making its precision drop to around 40% when HHalign stays around 60%. The two other pairs are AL00050021 and AL00052441 with respectively 14.61% and 15.38% sequence identity allowing HHalign to correctly align 60% of the target alignment. On AL00052441, PPalign correctly aligns more than 50% of the target alignment but the main difference comes here again from the precision (0.58 vs 0.81). Results on AL00050021 are clearly in favour of HHalign with an *F*_1_ score of 0.6 compared to 0.4 for PPalign and can be explained by the extremely gappy MSAs used to build the models (more than 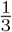 positions in the reference alignment were trimmed).

Interestingly, PPalign without coupling score (PPalign positional) achieves an *F*_1_ score comparable to HHalign (0.580 vs 0.578) despite a poor handling of gaps by Potts models as opposed to pHMMs. Besides, while PPalign’s alignment is most of the time better with the coupling score, 2 sequence pairs were yet significantly better aligned by PPalign positional than by PPalign with couplings: on already discussed AL10050875, where it improves a bit the poor quality of the alignment by PPalign, but also on AL00089447 (12.93% sequence identity) where it improves over the improvement of HHalign on PPalign.

## Discussion

Although the problem is very likely to be NP-hard since the threading problem is NP-hard [33], these experiments demonstrate that PPalign yields optimal Potts to Potts alignments up to a precision *ϵ* in tractable time. These results have to be confirmed on bigger instances. For now, experimentation is limited by memory handling in CCMpredPy, which is currently the only inference method offering the features we require to infer comparable Potts models, but the current implementation of CCMpred [25] shows that this type of inference can be optimized to handle significantly larger models. This should enable us to test larger alignments in the future. Based on our experimentation, we expect these alignments to be also tractable. This is surprising with respect to the NP-complete nature of the problem, but it seems that alignments of Potts models are not the hardest instances when they properly represent homologous proteins. We think that this depends yet on the choice of the parameters shaping the inference of Potts models and the similarity of the models to align: these questions deserve further studies to better understand the application scope of this method.

Regarding alignment quality, our results for the alignment of Potts models inferred using a pseudo-likelihood method designed for co-evolution prediction purposes are overall better than for the alignment of pHMMs by HHalign, with significant examples demonstrating how taking couplings into account can improve the alignment of remote homologous proteins, especially for lowest similarity alignments. There is still room for improvement in our method. We have noticed a tendency to align too many positions that can be corrected and our worst score with respect to HHalign is associated with very gappy train MSAs, indicating that augmenting Potts models with an appropriate gap handling strategy would undoubtedly improve our results. Above all, it is worth noting that PPalign positional finds sometimes a better alignment than PPalign, coupling matrices bringing more noise than assistance in these cases. To get better alignments, the priority is now is to work on more robust inference of Potts models, to make them more comparable and informative for homology search despite the relatively small size of training samples. We proposed here some ideas towards the inference of more canonical Potts models, with only the necessary couplings, as well as some post-processing steps, notably to smooth weights by simulated uniform pseudocounts. We are now searching for an efficient Potts model inference method that can be geared towards canonicity, providing the possibility to add pseudo-counts on the single and double amino acid counts – thus excluding methods based on pseudo-likelihood maximization – and would extend Potts models with an appropriate gap handling strategy.

## Conclusion

While Potts models have been successfully used for contact prediction and other tasks on protein sequences, using coevolutionary information captured by direct coupling analysis to improve homology search by sequence alignment seems promising, but challenging. The main computational bottleneck is the hardness of alignments involving Potts models.

We presented here PPalign, our method for Potts model to Potts model alignment based on the introduction of an Integer Linear Programming formulation of the problem with an implementation relying on an efficient solver able to yield the optimal solution in tractable time. This initiates a new approach for remote homology search by alignment of Potts models inferred from close homologs, similarly to HHalign with the alignment of pHMMs but with the addition of long distance sequence correlations reflecting the 3D structure of proteins. In this approach, Potts models need to be comparable. As a basic principle for building canonical Potts models, we proposed to infer models with as much weight as possible on the positional parameters and to add only necessary weight on pairwise couplings. We also proposed a scheme for lessening the effects of small sample variations on the Potts model’s parameters.

To experimentally assess the feasibility and interest of the approach, we carefully selected a set of non-redundant reference pairwise alignments with low sequence identity and with enough close homologs for each aligned sequence to infer a Potts models. We carried out rigorous experimentation with a strict separation of data used to train hyperparameters of the method and data used to test its performances. Results on test alignments confirm that Potts models can be aligned in reasonable time (1′37″ in average) and that taking into account direct coupling information can improve sequence alignments, especially for remote homologs with lowest sequence identity.

Our experiments suggest that new research on the inference of Potts models could improve their usefulness for homology search. The approach would undoubtedly benefit from extending to Potts models the insertion/deletions modeling capacities as well as the efficient pseudocount schemes of pHMMs. Maybe a more difficult issue is to have guarantees on a canonical form or at least some robustness of inferred Potts models to make them more comparable. We hope that PPalign’s efficiency and optimality will help to perform unbiased investigations in these directions.

## Acknowledgements

We would like to warmly thank Inken Wohlers for providing us with her code, Mathilde Carpentier for providing helpful scripts for alignment assessment and data since SISYPHUS database is temporarily unavailable, and the GenOuest Bioinformatics platform for providing computing resources.

## Availability of data and materials

Software and data are available here: https://www-dyliss.irisa.fr/ppalign

## Authors’ contributions

HT and FC devised the project and the main conceptual ideas. HT developed the theoretical formalism, carried out the implementation, built the benchmark and performed the experimentation. HT drafted the manuscript. HT and FC contributed to the final manuscript.

## Abbreviations

MSA: multiple sequence alignment
pHMM: profile Hidden Markov Model
ILP: Integer Linear Programming
UB: upper bound
LB: lower bound

